# SNPs in genes related to the repair of damage to DNA in clinical isolates of *M. tuberculosis*: a transversal and longitudinal approach

**DOI:** 10.1101/2023.12.05.570269

**Authors:** DE. Pérez-Martínez, R Zenteno-Cuevas

## Abstract

The presence of SNPs in genes related to DNA damage repair in *M. tuberculosis* can trigger hypermutagenic phenotypes with a higher probability of generating drug resistance. The aim of this research was to compare the presence of SNPs in genes related to DNA damage repair between sensitive and DR isolates, as well as to describe the dynamics in the presence of SNPs in *M. tuberculosis* isolated from recently diagnosed TB patients of the state of Veracruz, Mexico. The presence of SNPs in the coding regions of 65 genes related to DNA damage repair was analyzed. Eighty-six isolates from 67 patients from central Veracruz state, Mexico, were sequenced. The results showed several SNPs in 14 genes that were only present in drug-resistant genomes. In addition, by following of 15 patients, it was possible to describe three different dynamics of appearance and evolution of non-synonymous SNPs in genes related to DNA damage repair: 1) constant fixed SNPs, 2) population substitution, and 3) gain of fixed SNPs. Further research is required to discern the biological significance of each of these pathways and their utility as markers of DR or for treatment prognosis.

## Introduction

With more than 10 million new cases and 1.6 million deaths in 2021, tuberculosis (TB) remains one of the most important infectious diseases worldwide [1]. The increase in drug-resistant (DR) strains of *M. tuberculosis*, the greater susceptibility of hosts with diabetes mellitus and HIV/AIDS to contract the active infection, and the decrease in preventive and diagnostic activities by health systems in the face of the recent SARS-Cov2 pandemic, are the elements that currently influencing the severity of this pandemic [2].

SNPs are the main source of variation in *M. tuberculosis* [3] and their occurrence is influenced by drug treatment, host environment[4], and oxidative stress in the bacterium [5]. The high polymorphic presence in genes related to the DNA damage repair system (GRDDR) in *M. tuberculosis* has gained interest in recent years [6,7]. GRDDRs play a key role in genomic integrity and diversification, and thus in drug resistance [8,9]. Given this, it has been documented that the presence of SNPs in some GRDDRs can trigger hypermutagenic phenotypes that are more likely to generate DR [10,11] or conditions that can limit the survival of the bacterium [12].

Although research focused on the DNA damage repair system in *M. tuberculosis* has identified 65 GRDDRs, as well as the involvement of some of these in DR acquisition pathways [6], the polymorphic diversity of these genes, their usefulness as DR-TB markers, and the dynamics of SNP generation in GRDDRs during anti-TB treatment are still unknown. The objective of this research is therefore to compare the presence of SNPs in GRDDR between sensitive and DR isolates, as well as to describe the dynamics in the presence of SNPs in *M. tuberculosis* isolated from recently diagnosed TB patients of the state of Veracruz, Mexico.

## Materials and methods

### Collection and processing of isolates

Collection of clinical isolates of *M. tuberculosis* was performed by personnel from the State Health Services of Veracruz, Mexico, from January 2021 to June 2022. The sample size was based on convenience and depended directly on the number of patients diagnosed with TB by the health jurisdictions No. V and VII of the state of Veracruz, which are located in the municipalities of Xalapa and Orizaba, respectively.

Isolate processing was performed by collecting 5-15 ml of sputum from patients diagnosed with TB before initiating anti-TB treatment, with subsequent sampling every four weeks. Isolates were processed by Petroff’s method[13] and seeded on a solid Löwenstein-Jensen medium for ±8 weeks. Genomic DNA was then extracted and purified following the previously described CTAB method[14]. DNA was quantified using a nanodrop (ThermoScientific, USA). Only samples with DNA concentration >20 ng/μl, and a purity of 1.8-2.0 in ratio of absorbance at 260/280, and 2.0-2.2 in ratio of absorbance at 260/230, were considered for sequencing. Genomes were sequenced by NextSeq (Illumina). Genome sequences are available under the bioproject number PRJNA1041872.

### Bioinformatic analysis of the genomes

Low-quality ends (<30) were removed from the sequences using Fastp [15], then Kraken V.2[16] and SeqTK V1.3[17] were used to filter reads belonging to the MTBC complex [18] and avoid false variants as a result of DNA contamination.

Variant calling was performed using the MTBseq [19] and Tbprofiler [20] pipelines, an automated and modular process specialized in the mycobacteria of the *M. tuberculosis* complex, capable of identifying variants associated with resistance and phylogeny. By default, the pipeline uses the genome of *M. tuberculosis* H37Rv (NC_000962.3) as the basis for reference mapping, variant calling, and annotation. As a quality control, the genomes were also analyzed by PhyReSE[21] (https://bioinf.fz-borstel.de/mchips/phyresse/), identifying non-fixed allelic variants in the *M. tuberculosis* genomes.

From the variant calling, variants located in the coding regions of the 65 GRDDRs were identified and selected (Supplementary Table 1). The final database was constituted with the SNPs identified, resistance profile, lineage of the bacterium, patient ID, and date of isolate collection.

To identify the SNPs in GRDDR phylogenetically related to each *M. tuberculosis* sublineage, a phylogenetic analysis was performed using only the non-synonymous SNPs located in these genes using the IQ-TREE software[22] (http://iqtree.cibiv.univie.ac.at/) with the default parameters for binary data. The consensus tree generated was visualized with iTOL[23] (https://itol.embl.de/). Confirmation of the SNPs phylogenetically related to sublineages was performed using the fixation index (Fts = 1), which signals that an SNP is present in a lineage/sublineage but is not present outside of it. The fixation index was calculated with the Genepop package for Rstudio [24].

### Statistical analysis

Differences between sensitive and DR isolates were analyzed using Fisher’s exact test with the program IBM SPSS V21[25] (95% confidence level), excluding the SNPs phylogenetically associated with the sublineages. Comparison between consecutive intakes from the same patient was performed descriptively, based on the SNPs phylogenetically related to the infecting sublineage in each case.

## Results

### Genomic resistance and identified *M. tuberculosis* lineages

Eighty-six isolates from 67 patients from the center of the state of Veracruz, Mexico, were sequenced. A total of 67.4 % of the isolates were sensitive, while 32.6 % showed SNPs related to drug resistance. Among the drug-resistant isolates, 57 % were Pre-MDR, 18 % Mono-resistant, 14 % Pre-XDR, and 10 % MDR. Using the classification of Coll et al[26], 12 sublineages were identified, mainly from the Euro-American clade (L4), with the most frequent being L4.1.1, L4.1.1.3, and L4.1.2.1. Moreover, 100 % of L4.1.1.3 isolates showed some level of DR (Table 1). A total of 31.4 % of the isolates were collected before initiation of the anti-TB treatment.

**Table 1.** Distribution of sublineages observed in the sample and drug sensitivity profile.

| Lineage | DR |  | Sensitive |  | Total |  |
| --- | --- | --- | --- | --- | --- | --- |
|  | (n=28) |  | (n=58) |  | (n=86) |  |
|  | n | % | n | % | n | % |
| 4.1.1 | 0 | 0.0 | 24 | 41.4 | 24 | 27.9 |
| 4.1.1.3 | 17 | 60.7 | 0 | 0.0 | 17 | 19.8 |
| 4.1.2.1 | 2 | 7.1 | 10 | 17.2 | 12 | 14.0 |
| 4.3.3 | 3 | 10.7 | 9 | 15.5 | 12 | 14.0 |
| 1.2.1 | 3 | 10.7 | 2 | 3.4 | 5 | 5.8 |
| 4.1 | 0 | 0.0 | 4 | 6.9 | 4 | 4.7 |
| 4.9 | 2 | 7.1 | 1 | 1.7 | 3 | 3.5 |
| 4.3.4.2 | 0 | 0.0 | 3 | 5.2 | 3 | 3.5 |
| 4.8 | 1 | 3.6 | 1 | 1.7 | 2 | 2.3 |
| 4.3.4.1 | 0 | 0.0 | 2 | 3.4 | 2 | 2.3 |
| 4.3.4 | 0 | 0.0 | 1 | 1.7 | 1 | 1.2 |
| 4.4.1.2 | 0 | 0.0 | 1 | 1.7 | 1 | 1.2 |

DR: Drug-resistant isolate

### Variants in DNA repair genes

A total of 69 non-synonymous SNPs were identified and distributed among 38 GRDDRs in *M. tuberculosis*. The genes with the highest number of SNPs were *LigD* (5 sites), *DnaE2*, and *PolA* (4 sites). The only SNPs identified in *UvrD2* caused an early stop codon, while in *End* (*Nfo*) only a single deletion was observed. Although most of the SNPs identified were fixed in the genomes (allele frequency >95%), the *MutY, SSBb, RuvB, RecD*, and *AdnA* genes accounted for most of the SNPs with allelic variation, with a frequency of 5-94.9%.

#### Fixed variants associated with sublineages

The phylogenetic analysis of non-synonymous SNPs in GRDDR enabled the identification of 17 fixed SNPs related to the different sublineages that made up the sample (Table 2).

**Table 2.**
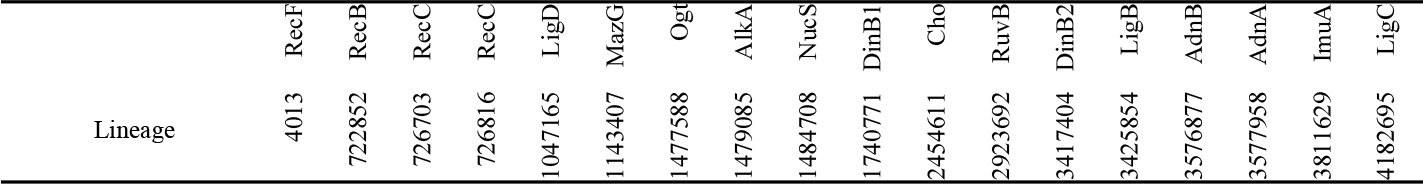
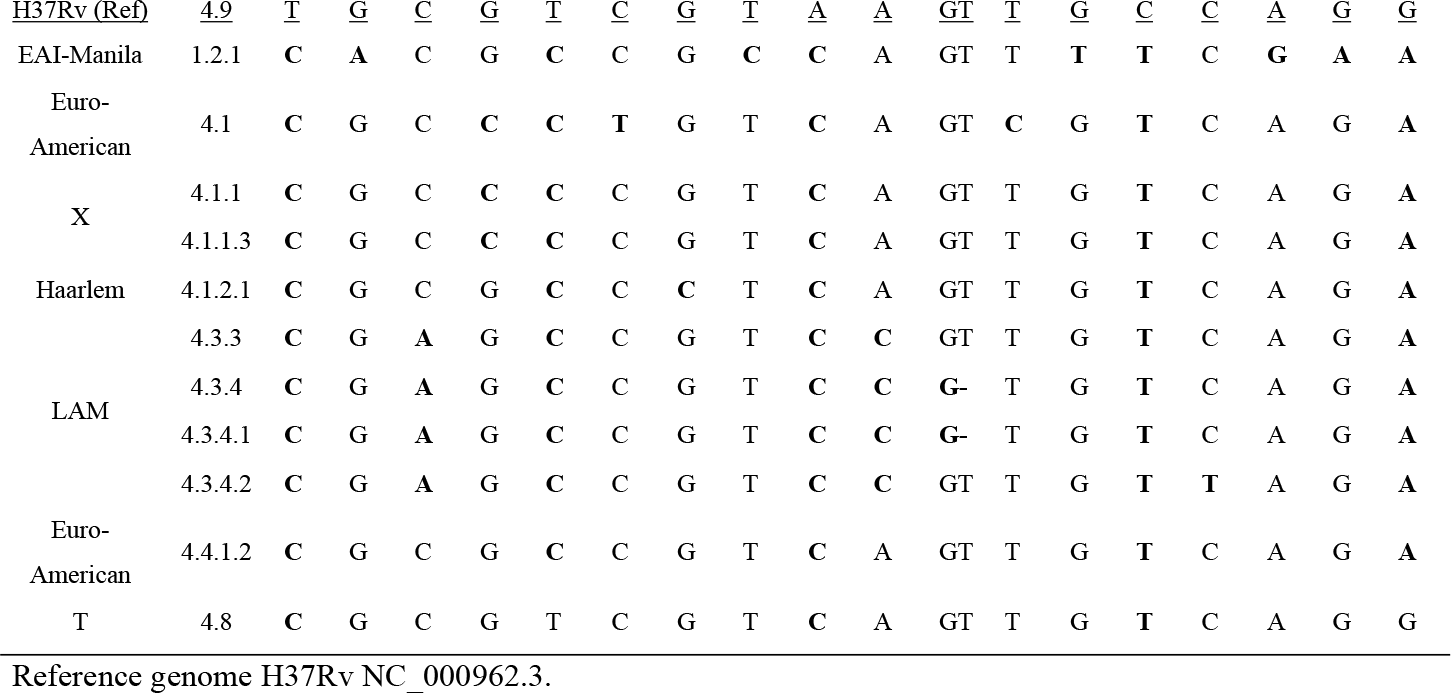
SNPs in DNA damage repair-related genes phylogenetically related to the sample sublineages.

#### Fixed variants associated with DR isolates

For most of the sublineages, some fixed SNPs were identified in GRDDR that differentiated between the DR and the sensitive genomes (Figure 1). For lineage X (L4.1.1. 3), two groups with distinctive SNPs were observed, the first of which was differentiated by the presence of the fixed SNPs *DinB2* (Asp172Asn) and *Prim-polC* (4180814 Leu303Val), where the presence of the fixed SNPs in *LigA* (3373792 Arg277Ser) differentiated between the Pre-MDR and pre-XDR isolates. The second group was distinguished by the presence of the SNPs *UvrB* (1837613 Asp180Gly) and *DnaE2* (3784559 Ala61Val). For the LAM lineage (L4.3.3), the SNPs *LigB* (3426223 Gly214Ser) and *AdnB* (3574941 Pro699Arg) were only observed in the Pre-MDR isolates. In EAI-Manila, the SNPs *UvrC* (1594594 Gly185Ser), *Ung* (3332159 Arg199Leu), *AdnA* (3579651 Arg184Cys), and *DnaE2* (3783802 Cys313Trp) were only observed in Pre-MDR genomes. In the T lineage (L4.8), the presence of *RNaseH1* (2502708, Ser11Ala) differentiated a single mono-resistant isolate. For the Haarlem isolates (L4.1.2.1), the SNPs *Fpg2* (1054138 Val125Ala) and *Dut* (3013784 Val122Leu) coincided in the DR genomes. It should be noted that no DR-related SNPs were observed for the H37Rv lineage (L4.9).

**Fig 1.**
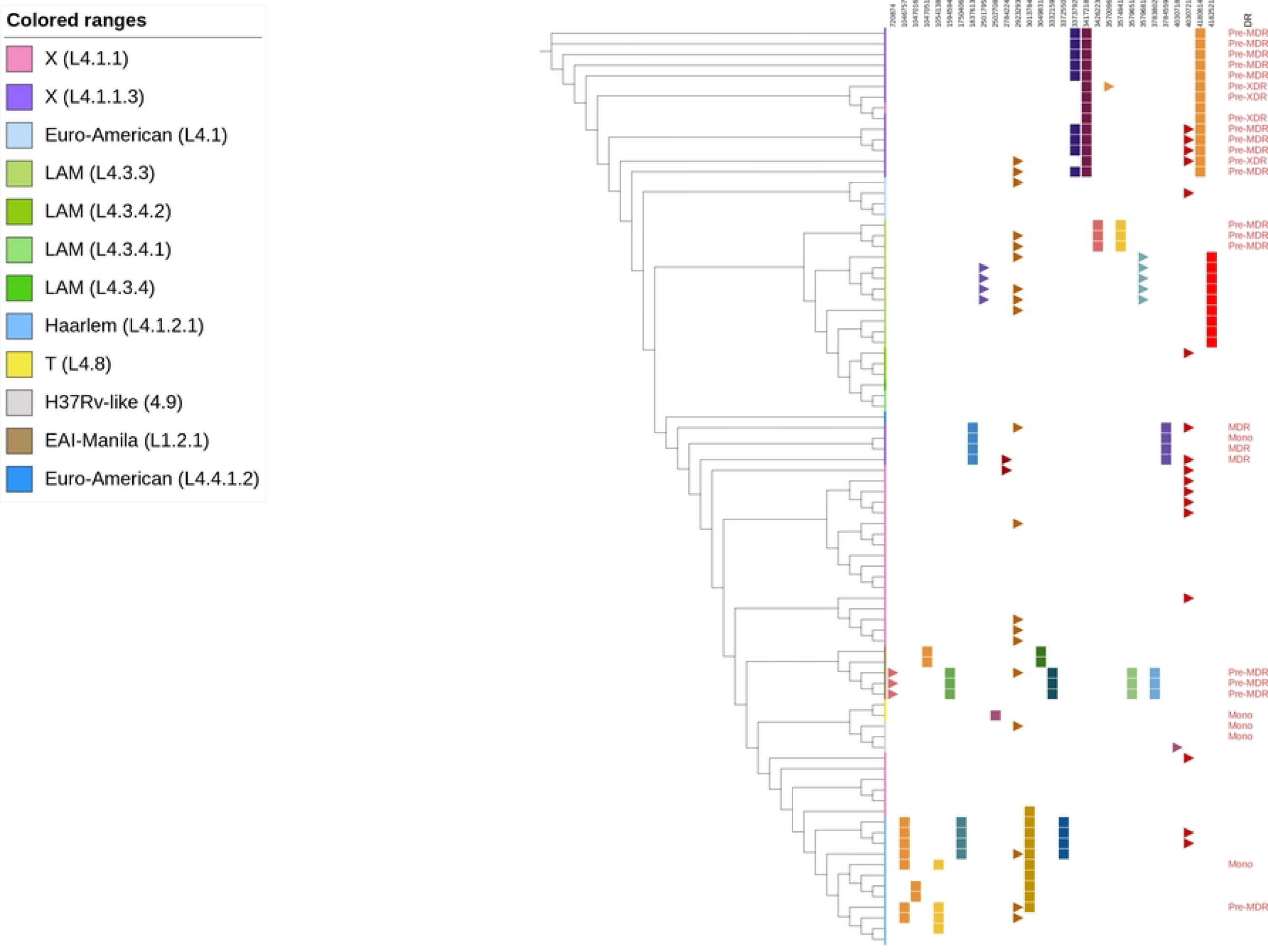
Fixed and variable SNPs in genes related to DNA damage repair according to lineage and drug resistance profile. DR: drug resistance profile. Mono: Mono-resistant. Squares: Fixed SNPs. Triangles: SNPs with allelic variation.

Notably, the *MutY* SNPs (4030721, Ala77Pro) showed allelic variation in some sensitive and DR isolates belonging to the L4 lineage, while the *RuvB* SNPs (2923293, Arg314Pro) also showed allelic variation in both the L4 and L1 genomes. Likewise, the SNPs *RNaseH1* (2501795, Thr315Met) and *AdnA* (3579681, Asp174Asn) showed allelic variation in sensitive L4.3.3 genomes, while *RecD* (720874, Arg287Gly) SNPs were variant in DR genomes of L1.2.1 (Figure 1).

Not considering SNPs related to sublineages (Table 2), the comparison between isolates showed significant differences in the presence of 12 SNPs in DR genomes and three SNPs present in sensitive isolates (Table 3). Among these, the presence of non-synonymous SNPs in the genes *DinB2* or *DnaE2* observed in 71.4% of the DR genomes was notable. On the other hand, the presence of non-synonymous SNPs in the *Nei1* gene was observed in 31% of the sensitive isolates.

**Table 3.**
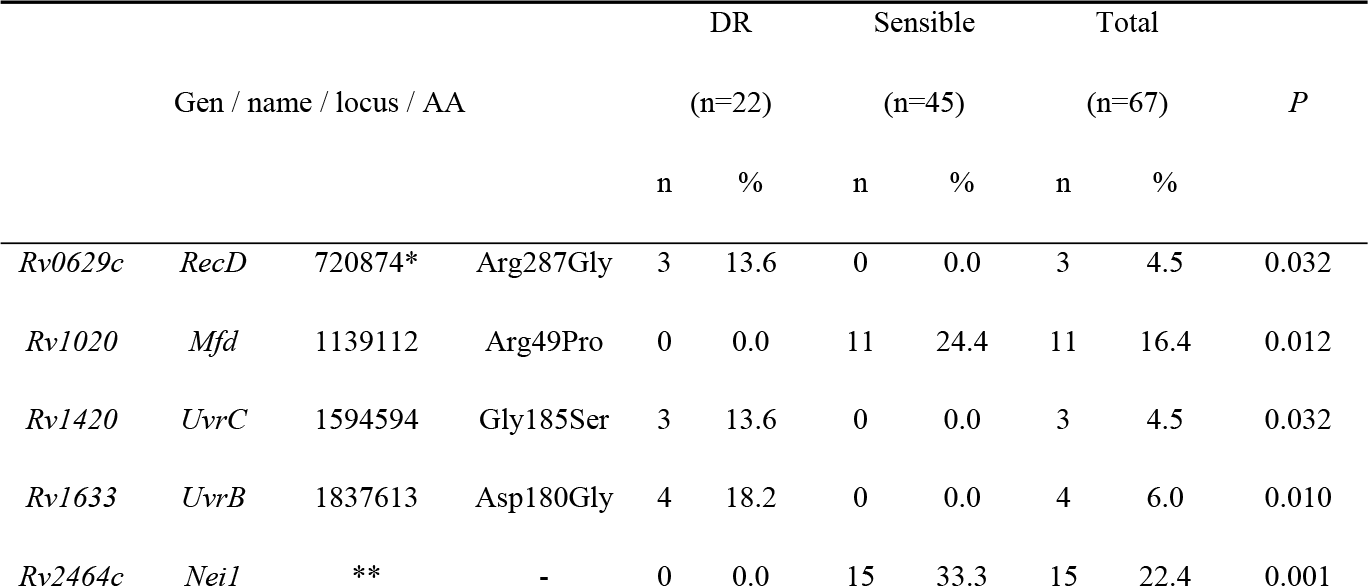
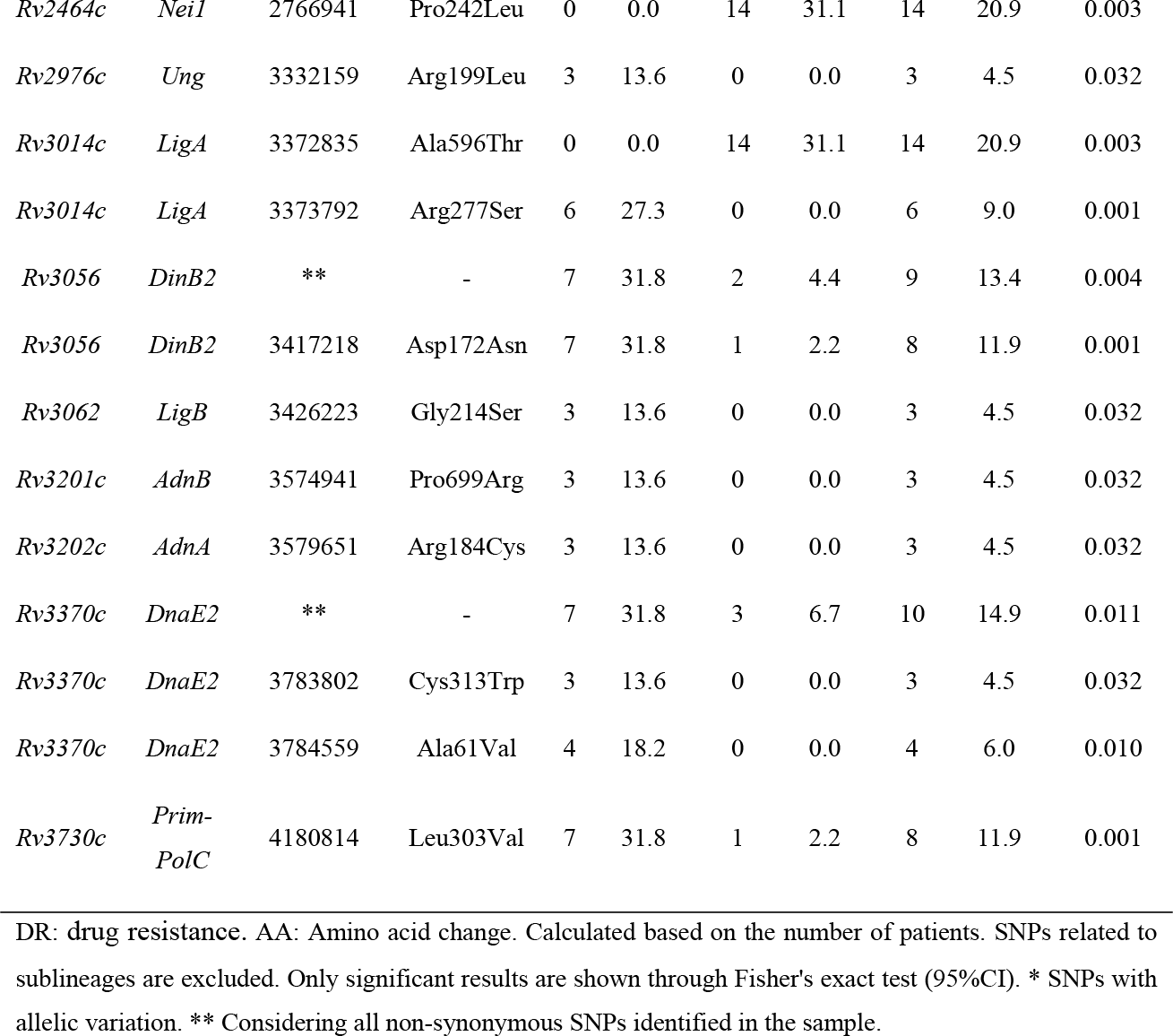
Genes and SNPs related to sensitive and drug-resistant genomes.

### Changes in the presence of SNPs in GRDDR during anti-TB treatment

The longitudinal analysis allowed us to follow the changes in GRDDRs in *M. tuberculosis* before and after the standardized anti-TB treatment (R/Z/E/P). Through a follow-up of 15 patients, it was possible to describe three different dynamics of appearance and evolution of non-synonymous SNPs in GRDDR (Figure 2).

**Fig 2.**
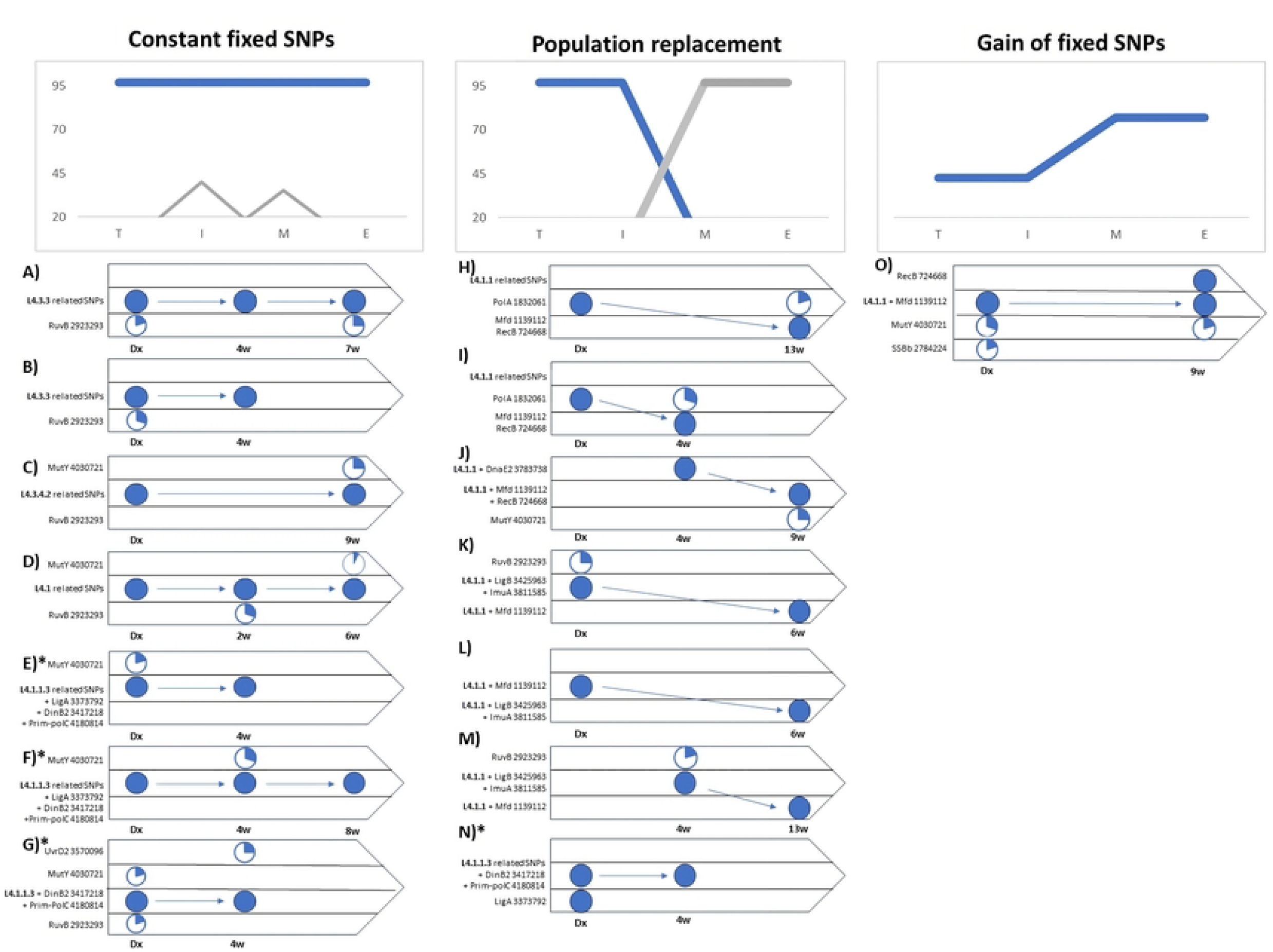
Dynamics of SNPs in genes related to DNA damage repair during treatment. **W:** weeks. Arrow: changes over time. Circles: percentage of allele frequency in the bacterial population.

1. Constant fixed SNPs: these are related to the sublineages and remain constant between the different isolates of a patient (Figure 2 A-G). Although allelic diversification was observed in some loci, it was sporadic and they therefore did not become fixed as dominant SNPs in the infecting population. This dynamic was mainly observed in sensitive genomes with the sublineages L4.1, 4.3.3, and 4.3.4.2. The constancy of fixed SNPs between consecutive isolates was also observed in three patients with DR-TB caused by L4.1.1.3.
2. Population substitution of SNPs: these differ between pre- and post-treatment intakes, suggesting population substitution; i.e., the initial population decreases and disappears, while another population increases until it becomes the dominant population. This type of diversification was observed in isolates with the lineages L4.1.1 and L4.1.1.3 (Figure 2 H-N).
3. Gain of fixed SNPs: In addition to sublineage-related SNPs, genomes are gaining fixed SNPs in their GRDDRs in their consecutive isolates. This was observed in a single patient L4.1.1 (Figure 2 O).

## Discussion

The present study represents the first approach focused on the allelic diversity of GRDDRs in *M. tuberculosis* circulating in affected population. The analysis allowed us to identify some SNPs present only in DR isolates, as well as to observe allelic variation in genes recently related to hypermutagenic TB phenotypes. In addition, analysis of consecutive samples allowed us to describe, for the first time, how the presence of SNPs in GRDDRs can change between pre- and post-treatment isolates, following three different dynamics. It is worth mentioning that the small number of samples limited the analysis to a descriptive scope.

SNPs observed only in DR isolates were located in the low-fidelity polymerases *DinB2* [27] and *DnaE2* [28], as well as the polymerase *Prim*-*polC* [29], which reportedly participate in induced mutagenesis in *M. tuberculosis*; the ligases *LigA* and *LigB* [6]. SNPs were also observed in the *Uvr*-*BC* genes involved in DNA damage recognition and initiation of repair by nucleotide excision[6], in which the presence of mutations has been associated with failure of the repair pathway and consequently increased mutations and drug resistance [11]. In the same sense, they were identified in the *Adn-AB* dimer, a participant in the initiation of DNA cleavage repair by *RecR*-dependent homologous recombination[6]. The uracil-specific glycosylase *Ung* [30], is a gene in which the presence of SNPs is linked to the acquisition of DR in the Haarlem lineage[31]. Also were observerd the ribonucleotide glycosylase *RNaseH1*[6]; the non-functional *MutM*/*Fpg* homolog, *Fpg2* [6]; as well as *Dut*, an enzyme involved in the nucleotide pool sanitization of dUTP and dCTP [6]. Although the presence of differential SNPs in DR isolates may be due to selective treatment pressure, some compensatory mechanism, or related to an increased adaptability of the bacterium, their implications for disease control, and their utility as surrogate markers of DR-TB are unknown.

Although none of the SNPs with low allele frequency showed significant associations with DR, the allelic variation in *MutY* (4030721, *Ala*77Pro) and *RuvB* (2923293, Arg314Pro) present in different sublineages could suggest a homoplasy event. Although *RuvB* is a protein involved in the resolution of branched DNA structures[32], our previous observations suggest that the occurrence of SNPs in *RuvB* is more frequent in the presence of DMT2 in the host [33]. On the other hand, recent research has shown that some mutations in *MutY* abrogate its functions generating a hypermutagenic phenotype, with increased survival and rapid acquisition of resistance mutations for rifampicin and ciprofloxacin [11,34]. On the other hand, the presence of SNPs in *Mfd* and *RecB* observed in sensitive genomes of L4.1.1 with >9 weeks of anti-TB treatment could reflect positive selection for anti-TB drugs, while the absence of SNPs related to drug resistance in these isolates could indicate that the SNPs in *Mfd* and *RecB* are related to persistence or tolerance pathways of *M. tuberculosis*.

Analysis of consecutive isolates showed that the presence of SNPs in GRDDR can fluctuate during the anti-TB treatment. Consequently, it was possible to elucidate three distinct dynamics: 1) constant fixed SNPs, 2) population substitution, and 3) gain of fixed SNPs. Although SNP gain and population substitution have been previously documented as adaptive pathways of *M. tuberculosis*[35–37], our findings represent the first report of this perspective of analysis using only SNPs in GRDDR, highlighting the importance of these genes, and their diversification, in the early months of anti-TB treatment.

The constant presence of fixed SNPs among serial isolates could imply a functional GRDDR SNP configuration, both for infection and for coping with the anti-TB treatment. Although allelic variation in some other genes was observed in this configuration, it was transient and was not observed to be fixed in the dominant population. This is consistent with other studies documenting long-term infections, with no changes in the presence of fixed SNPs in populations of *M. tuberculosis*[37,38].

On the other hand, population substitution could indicate that the configuration of SNPs in GRDDR observed in the initial strain was unable to withstand the damage generated by drug therapy, such that the initial population is no longer observed as predominant and is replaced by bacteria with a different configuration of SNPs[36,39]. We can also theorize that the event of population substitution will depend on the diversity of bacterial subpopulations present in the host. In this regard, observations in some *M. tuberculosis* infections with the L4.1.1 sublineage suggest that genomes with non-synonymous SNPs in *RecB* and *Mfd* were better able to tolerate drugs, even if they were not the dominant populations identified at the onset of symptomatology and diagnosis by health services.

The gain of fixed SNPs in GRDDR was the least observed dynamic in the sample and its presence opens new questions about its origin. Although the difference in SNPs observed between consecutive isolates could indicate the selective pressure of pre-existing bacterial subpopulations in the face of drug treatment, it is also possible that these SNPs were generated *de novo* in the dominant population. The fact that the only case where a gain of fixed SNPs was observed was a sensitive TB (L4.1.1) could indicate that the diversification of GRDDRs is one of the prior routes for DR acquisition, the origin of which could be related to the time of drug exposure and host-specific factors.

With regard to the diversity of *M. tuberculosis* observed in the state of Veracruz, the prevalence of Euro-American (L4) lineages of TB in Mexico has been documented since the last decade[40]. However, the incidence of lineage X (L4.1.1 and L4.1.1.3) increased in the region, from ∼13 % (2012-2021)[41,42], to the 42 % reported in this study. This is of concern given the high frequency of drug resistance in lineage L4.1.1.3 previously identified in this [43,44] and other regions[45].

## Conclusion

Analysis of *M. tuberculosis* isolates showed SNP variations in GRDDR between drug-resistant and drug-sensitive isolates. Furthermore, a comparison of pre- and post-treatment isolates indicates that the diversification of SNPs in GRDDR could have three distinct dynamics: 1) constant fixed SNPs, 2) population substitution, and 3) gain of fixed SNPs. However, further research is required to discern the biological significance of each of these pathways and their utility as markers of DR or for treatment prognosis.

## Funding

This project financed with resources from CONACYT-Basic Sciences Fund: Influence of type 2 diabetes mellitus in the development of mutations associated with multidrug resistance in Tuberculosis, A1-S-22956. The doctoral student was granted scholarships by CONACYT, call “Becas Nacional (Tradicional) 2019-02” with registration number (“CVU”) 411155.

## Acknowledgments

We thank to Veracruz, Canales-Velásquez G, and Arroyo C. of the Mycobacteria program of the Health secretary of for their invaluable support in the collection of clinical isolates.

## Contribution roles

DE-PM work in the conceptualization, data curation, formal analysis, validation, writing draft and review and editing. R-ZC work in the conceptualization, formal analysis, supervision, funding acquisition, writing draft and review and editing.

## Notes

### Competing Interest Statement

The authors have declared no competing interest.

